# Towards Translational Sleep Staging: A Cross-Species Deep-Learning Model for Rodent and Human EEG

**DOI:** 10.64898/2026.02.25.707936

**Authors:** Bartlomiej Chybowski, Alfredo Gonzalez-Sulser, Javier Escudero

**Affiliations:** Muir Maxwell Epilepsy Centre, University of Edinburgh, Street, Postcode, Edinburgh, Scotland; School of Medicine, Deanery of Clinical Sciences, University of Edinburgh, 50 Little France Crescent, EH16 4TJ, Edinburgh, Scotland; School of Engineering, Institute for Imaging, Data and Communications, University of Edinburgh, University of Edinburgh, Alexander Graham Bell Building, Thomas Bayes Road, EH9 3FG, Edinburgh, Scotland; School of Medicine, Centre for Discovery Brain Sciences, University of Edinburgh, 1 George Square, EH8 9JZ, Edinburgh, Scotland

**Keywords:** Electroencephalography, Deep Learning, Cross-species translation, Sleep staging, Rodent models, Polysomnography

## Abstract

**Study Objectives:** Automated sleep staging underpins clinical sleep assessment and translational neuroscience, yet most data analyses work addresses human and animal data separately. We tested whether a seizure-oriented machine learning framework can be repurposed for three-state sleep staging in humans and rats, and whether models trained solely on rodent data can be applied directly to human recordings using an explicit cross-species montage.

**Methods:** We used the PySeizure, a standardised EEG preprocessing and seizure-detection framework, together with TinySleepNet as the core classifier. Models were trained and evaluated on the Sleep-EDF expanded Sleep Cassette subset (three classes: wake, non-rapid eye movement sleep, rapid eye movement sleep), then applied without fine-tuning to the Sleep Telemetry subset. The same pipeline was used on a SYNGAP1 rat dataset with analogous three-state labels. A novel human–rat electroencephalography montage mapped rat electrodes to putative human scalp homologues, enabling direct application of rat-trained models to Sleep Cassette.

**Results:** Within Sleep Cassette, the accuracy in three-stage sleep classification was 0.95. Applying this model directly to Sleep Telemetry yielded an accuracy of 0.89. On the rodent dataset, accuracy was 0.78. When the rat-trained model was applied directly to Sleep Cassette, accuracy was 0.68.

**Conclusions:** A single deep learning pipeline can support robust three-state sleep staging in humans and rodents and retains meaningful performance under both human cross-subset and rat-to-human transfer without any retraining or fine-tuning. The rat-trained model’s above-chance performance on human data, achieved without human training samples, shows that rodent-derived representations can contribute directly to human sleep staging when constrained by an anatomically informed montage, linking preclinical rodent recordings and clinical human sleep studies.

## Introduction

Sleep staging, the classification of polysomnography recordings into canonical vigilance states, is fundamental to clinical sleep medicine and basic sleep research [14, 6]. In routine practice, expert scorers segment overnight recordings into fixed-length epochs, most commonly 30 seconds, and assign each epoch to wake, non-rapid eye movement (NREM) sleep, or rapid eye movement (REM) sleep according to established criteria such as the Rechtschaffen and Kales rules or the American Academy of Sleep Medicine (AASM) scoring manual [10, 17]. Although manual scoring provides the de facto reference standard, it is time-consuming, resource-intensive, and subject to inter- and intra-scorer variability [5], motivating the development of accurate, reproducible, and scalable automated approaches.

Over the past decade, deep learning (DL), a subfield of machine learning (ML), has become the dominant paradigm for automatic sleep staging from electroencephalography (EEG) and other signals used in polysomnography (PSG). Models based on convolutional neural network (CNN) and recurrent neural network (RNN) operating on raw or minimally processed EEG have achieved performance that approaches human inter-scorer agreement in several benchmark datasets [21, 18, 15]. Architectures specifically designed for sleep staging, such as TinySleepNet [19], demonstrate that compact DL models can attain high accuracy using a single EEG channel, enabling deployment in wearable and resource-constrained settings. More recent work in artificial intelligence (AI) for sleep has further explored end-to-end models for single- and multi-channel EEG, as well as attention-based and explainable artificial intelligence (XAI)-oriented architectures, to improve robustness and interpretability [13, 16].

Nevertheless, an important gap remains. Most existing ML-based sleep staging methods have been developed separately for human or for rodent data, typically using different label schemes, preprocessing pipelines, and architectures. There is a lack of systematic, data-driven approaches that deploy a single modern ML model with harmonised three-state labels (wake, NREM, REM) across both humans and animal models within one unified training and evaluation framework, and explicit assessments of model transfer across species are rare or absent. In parallel, there is growing interest in explicitly translational and cross-species approaches to sleep, in which mechanistic insights from rodent models are related to human sleep physiology and pathology [1, 9]. Comparative neuroanatomical and functional studies indicate a conserved organisation of cortical networks across mammals, including homologous frontal and parietal regions that contribute to sleep oscillations and vigilance regulation in both rodents and humans [8, 1]. These correspondences suggest that, with an appropriate mapping between rodent electrode locations and human scalp sites, it should be possible to assess whether ML models trained exclusively on rodent EEG can provide non-trivial sleep staging performance when applied directly to human recordings. To our knowledge, such cross-species transfer of a modern DL sleep staging model trained only on rodent data has not yet been systematically evaluated. Establishing this form of transfer would provide a quantitative bridge between preclinical models and clinical sleep phenotypes, supporting genuinely translational analyses of sleep across species.

In this study, we address these gaps by extending PySeizure [3], a standardised EEG preprocessing and seizure-detection framework, with automated sleep-stage detection in humans and rats, integrating TinySleepNet [19] as the core DL model. PySeizure was originally developed for epilepsy research, where sleep and circadian organisation are known to modulate seizure occurrence; in the present work, we therefore focus on the sleep-staging component, while retaining the same underlying infrastructure. Classification is restricted to three vigilance states (wake, NREM, REM) across species for two reasons. First, the available rat dataset is annotated only with wake, NREM, and REM labels. Second, although finer-grained substaging of NREM in rodents has recently been proposed [4, 22], these schemes are not yet universally adopted, which complicates direct alignment with standard human N1–N3 scoring. Based on known anatomical and functional correspondences between rodent and human cortical regions [20], we then define a cross-species EEG montage that maps rat recording sites to putative human scalp homologues. First, we train and evaluate the PySeizure-TinySleepNet pipeline on the sleep-edf dataset extended (SleepEDFx) sleep cassette (SC) subset, treating it as a human benchmark for three-state sleep staging. Second, we perform a cross-subset generalisation experiment by applying the models, without any fine-tuning or calibration, to the SleepEDFx sleep telemetry (ST) subset, thereby probing domain shift between cassette-based and telemetry-based human recordings under the same three-state label scheme. Third, we train and evaluate the same pipeline on a rat EEG dataset (SYNGAP1 rodent dataset (SRD)) with wake, NREM, and REM annotations. Finally, we conduct a cross-species transfer experiment in which models trained exclusively on rat EEG are applied directly to the SC subset, again without using any human data for training, model selection, or calibration. Across these experiments, we aim to provide a proof-of-concept for a unified, cross-species approach to automated sleep staging and to investigate the extent to which representations learned in a rodent model can be translated to human sleep.

## Methods

### Datasets

#### Human data: Sleep-edf dataset extended

Human PSG data were obtained from the SleepEDFx [12]. We used the canonical PhysioNet release of SleepEDFx, in which all signals are resampled to 100 Hz [7]. For this study, all EEG channels used as model inputs were further resampled to 256 Hz to ensure that human and rodent data shared the same sampling frequency. Each recording typically spans about 7–10 hours from lights-off to final awakening. We used its two principal subsets:

- **Sleep cassette:** 153 overnight PSG recordings from 78 healthy subjects, recorded mostly at home to study age effects on sleep.
- **Sleep telemetry:** 44 overnight PSG recordings from 22 subjects, recorded in hospital to investigate the effects of temazepam versus placebo, typically two nights per subject (drug and control).

Sleep stages in SleepEDFx are annotated every 30 seconds according to the Rechtschaffen and Kales rules. The original hypnograms include wake, NREM stages 1–4, REM, movement time, and, in some recordings, unscored or artefactual epochs. To harmonise the label space with the rodent annotations, we mapped the original human labels to a three-state scheme: wake, NREM, and REM. All NREM stages (N1–N4) were merged into a single NREM class. Any epochs not assigned to one of these three states were excluded from training and evaluation.

#### Rodent data: SYNGAP1 rodent dataset

Rodent data were obtained from the SRD, which contains EEG recordings from rats that model SYNGAP1 haploinsufficiency, generated at the Simons Initiative for the Developing Brain at the University of Edinburgh by the laboratory of Dr Gonzalez-Sulser [2, 11]. We used recordings from nine animals, acquired with the H16-EEG-NeuroNexus Grid system using a 14-channel cortical montage plus 2 electromyography (EMG) channels at an original sampling frequency of 250.4 Hz. All channels were resampled to 256 Hz for consistency with the human data.

Each rat contributed at least one long-term recording (24 hours), and three had continuous recordings of approximately 48 hours. Vigilance states were scored manually into three classes (wake, NREM, REM) in 5-second epochs, and these native epochs and labels were used directly without temporal aggregation. All available recordings from these nine rats were included in the analysis.

#### Preprocessing

All preprocessing was performed within the PySeizure framework (Section 2.4) [3]. For both human and rodent data, we followed the preprocessing steps described in the original PySeizure work, with one different step applied in this study: per-subject z-score normalisation.

For SleepEDFx, we used the EEG signals as provided in the SleepEDFx, resampled them from 100 Hz to 256 Hz, segmented them into 30-second epochs aligned with the provided hypnograms, and applied the label mapping to wake, NREM, or REM as described above. Epochs annotated with any other label were excluded. For each subject and each channel, we then applied per-subject z-score normalisation by subtracting the mean and dividing by the standard deviation (STD) computed over the full recording.

For the SRD SYNGAP1 rats, all 14 channels were resampled from 250.4 Hz to 256 Hz. We used the manually scored 5-second epochs with wake, NREM, and REM labels and automatically excluded from training any epochs flagged as artefactual either in the original annotations or by the PySeizure artefact routines. For each rat and each channel, we also applied per-subject z-score normalisation.

No additional hand-crafted features were computed; TinySleepNet operated directly on these preprocessed, normalised EEG time series.

#### TinySleepNet and architectural modification

TinySleepNet is a compact DL model for automatic sleep staging from raw single-channel EEG [19]. In its original form, it applies a series of convolutional layers to a fixed-length input epoch (typically 30 seconds), followed by temporal aggregation and a final classifier that outputs sleep stage probabilities.

In this work, TinySleepNet served as the core classifier inside PySeizure for both human and rodent data. We retained the default configuration from the original work for the convolutional feature extractor (kernel sizes, numbers of filters, activation functions) and modified only the final part of the network so that it no longer depended on a specific epoch length.

In the original implementation, the final classification layer expects a fixed-size feature vector determined by the epoch duration and sampling frequency. We removed this dependency by introducing a global average pooling operation over the last convolutional feature maps, followed by a fully connected layer whose input dimension depends only on the number of filters in the final convolutional layer. This makes the classifier independent of the input epoch length. After TinySleepNet has produced its final feature maps, these are averaged across time to obtain a fixed-length feature vector, regardless of the original epoch duration. This vector is then passed to a linear layer that outputs three logits corresponding to wake, NREM, and REM. This modification preserves the original representation learning of TinySleepNet while allowing the same model instance to process both 30-second human epochs and 5-second rodent epochs at 256 Hz. All models were trained with categorical cross-entropy over the three classes (wake, NREM, REM), without explicit class weighting.

#### PySeizure pipeline

PySeizure [3] is a Python-based signal processing and ML framework originally developed for seizure detection from multichannel EEG. It provides configuration-driven data loaders, reusable preprocessing modules, wrappers for DL models implemented in PyTorch, training utilities, and evaluation routines.

For this study, PySeizure was extended to support sleep staging as a native task across human and rodent data. The main adaptations were:

- support for epoch-based three-class labelling (wake, NREM, REM);
- dataset handlers for SleepEDFx (SC and ST) and SRD SYNGAP1 rats, including the label mapping to wake, NREM, and REM;
- integration of TinySleepNet, with the length-agnostic modification, as a PySeizure model component; and
- configuration profiles for human-only, rodent-only, and cross-species training and evaluation.

All preprocessing, model instantiation, and training/evaluation were executed via PySeizure configuration files. For complete transparency and to enable replication, the code is available in the GitHub repository associated with the original PySeizure paper [3].

#### Human–rat EEG mapping

To enable cross-species experiments, we constructed a human–rat montage that maps bipolar combinations of human scalp electrodes to corresponding bipolar pairs of rat cortical electrodes in the H16-EEG-NeuroNexus Grid montage (Table 1).

**Table 1.**
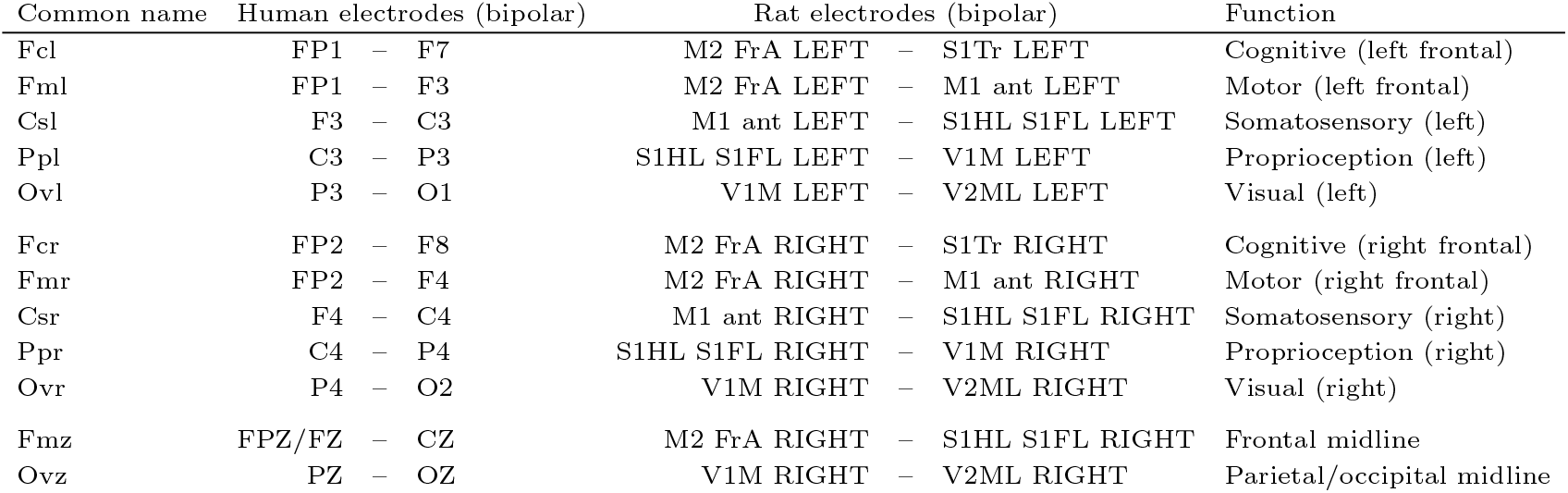
Human–rat bipolar EEG mapping used for cross-species experiments. Each custom channel is defined by a pair of human scalp electrodes and a corresponding pair of rat H16-EEG-NeuroNexus Grid electrodes, together with a descriptive functional label.

| Common name | Human electrodes (bipolar) | Rat electrodes (bipolar) | Function |
| --- | --- | --- | --- |
| Fcl | FP1 – F7 | M2 FrA LEFT – S1Tr LEFT | Cognitive (left frontal) |
| Fml | FP1 – F3 | M2 FrA LEFT – M1 ant LEFT | Motor (left frontal) |
| Csl | F3 – C3 | M1 ant LEFT – S1HL S1FL LEFT | Somatosensory (left) |
| Ppl | C3 – P3 | S1HL S1FL LEFT – V1M LEFT | Proprioception (left) |
| Ovl | P3 – O1 | V1M LEFT – V2ML LEFT | Visual (left) |
| Fcr | FP2 – F8 | M2 FrA RIGHT – S1Tr RIGHT | Cognitive (right frontal) |
| Fmr | FP2 – F4 | M2 FrA RIGHT – M1 ant RIGHT | Motor (right frontal) |
| Csr | F4 – C4 | M1 ant RIGHT – S1HL S1FL RIGHT | Somatosensory (right) |
| Ppr | C4 – P4 | S1HL S1FL RIGHT – V1M RIGHT | Proprioception (right) |
| Ovr | P4 – O2 | V1M RIGHT – V2ML RIGHT | Visual (right) |
| Fmz | FPZ/FZ – CZ | M2 FrA RIGHT – S1HL S1FL RIGHT | Frontal midline |
| Ovz | PZ – OZ | V1M RIGHT – V2ML RIGHT | Parietal/occipital midline |

The mapping was based on the available implantation sites and guided by comparative neuroanatomical principles describing conserved cortical organisation across mammals [20]. In particular, it was designed to pair anterior—posterior and lateral positions with similar putative functional roles, for example, frontal association and motor areas, primary somatosensory cortex, or visual cortex. Human left/right frontal, central, parietal, and occipital derivations were therefore matched to rat electrodes overlying frontal association/motor, somatosensory, and visual cortices in the corresponding hemisphere, while midline human derivations were mapped to right-hemisphere rat sites because the implant does not include true midline cortical electrodes. For each custom channel, we thus defined a pair of human electrodes combined as a bipolar derivation, a corresponding pair of rat H16-EEG-NeuroNexus Grid channels combined as an analogous bipolar derivation, and a short functional description. This design does not imply a one-to-one homology between individual electrodes, but rather an approximate alignment of major cortical territories and their dominant functional roles.

## Results

We evaluated the PySeizure–TinySleepNet pipeline in four configurations: (i) within-dataset evaluation on SC, (ii) cross-subset generalisation from SC to ST, (iii) within-dataset evaluation on the SRD dataset, and (iv) cross-species transfer from SRD rats to SC using the human–rat montage. All metrics were computed over all epochs and all three classes (wake, NREM, REM) jointly.

### Within-SC evaluation (SC → SC)

When trained and evaluated on SC, the model achieved an accuracy of 0.95, area under the receiver operating characteristic curve (ROC AUC) of 0.99, sensitivity of 0.95, and specificity of 0.98, indicating excellent three-state sleep staging performance under within-subset conditions.

### Cross-subset human generalisation (SC → ST)

Applying the same SC-trained model to ST without any adaptation resulted in a reduction in performance (accuracy 0.89, ROC AUC 0.96, sensitivity 0.88, specificity 0.92), but performance remained high, showing that the pipeline generalises well across differences in human acquisition context.

### Within-rodent evaluation (SRD → SRD)

On the SRD, the model reached an accuracy of 0.78, ROC AUC of 0.96, sensitivity of 0.77, and specificity of 0.91, demonstrating good within-species performance in rodents using the same architecture and training framework.

### Cross-species transfer (SRD → SC)

Finally, when the rat-trained model was applied directly to SC via the human–rat montage, performance decreased (accuracy 0.68, ROC AUC 0.75, sensitivity 0.68, and specificity 0.74), but remained clearly above chance, indicating that representations learned from rat cortical activity retain non-trivial predictive value for human sleep staging in the absence of any human training data.

## Discussion

In this study, we adapted the **PySeizure**, a standardised EEG preprocessing and seizure-detection framework originally developed for epilepsy research [3], together with **TinySleepNet** [19] as the core classifier to perform three-state sleep staging (wake, NREM, REM) across human and rodent EEG recordings and to test cross-species transfer using an anatomically informed human–rat montage. Here we summarise the empirical findings and discuss their translational and clinical relevance, as well as how they reflect the design principles of PySeizure.

### Translational and clinical implications

#### Human performance and deployment across recording contexts

Within the SC subset of SleepEDFx, the PySeizure– TinySleepNet pipeline achieved high performance (Table 2) under a three-state (wake, NREM, REM) labelling scheme.

**Table 2.** Summary of model performance across all experimental conditions, comprising evaluations on the sleep cassette (SC), sleep telemetry (ST), and SYNGAP1 rodent dataset (SRD) databases. Each result column denotes a paired training and evaluation setting. Performance is quantified using mean square error (MSE), area under the receiver operating characteristic curve (ROC AUC), accuracy, precision, area under the precision recall curve (PRAUC), sensitivity, and specificity, with all metrics computed jointly for the three sleep-stage classes: wake, NREM, and REM. The arrow symbol (→) denotes that models were trained on the dataset indicated before the arrow and evaluated on the dataset indicated after the arrow.

| Metric | SC → SC | SC → ST | SRD → SRD | SRD → SC |
| --- | --- | --- | --- | --- |
| MSE | 0.02221 | 0.05601 | 0.11070 | 0.15406 |
| ROC AUC | 0.99459 | 0.96177 | 0.95688 | 0.74517 |
| Accuracy | 0.95269 | 0.88716 | 0.77753 | 0.68261 |
| Precision | 0.86609 | 0.83804 | 0.67576 | 0.54157 |
| PRAUC | 0.85848 | 0.85397 | 0.45609 | 0.16819 |
| Sensitivity | 0.95269 | 0.88716 | 0.77753 | 0.68261 |
| Specificity | 0.98097 | 0.91916 | 0.91147 | 0.74013 |

This indicates that, when training and testing on data from the same acquisition protocol, the model can provide reliable automatic sleep staging that is consistent with levels typically reported for clinical and research applications. It is important to note, however, that unlike many previous studies on SleepEDFx that perform five-class staging (wake, N1, N2, N3/N4, REM), the present work uses a three-class label scheme that merges N1–N4 into a single NREM class. This simplification is necessary to harmonise human and rodent labels, but it also makes direct numerical comparisons to published five-class results imperfect, as the task considered here is somewhat easier. Within this three-class setting, our SC → SC performance lies at the upper end of previously reported single-channel methods.

When the same model, trained only on SC, was applied without any recalibration to the ST subset, performance decreased but remained high (Table 2). These results show that a single model trained on cassette-based recordings can retain useful accuracy on telemetry-based recordings, despite differences in recording setup and context. For clinical deployment, this suggests that models trained on relatively standard PSG collections could be applied to related in-hospital or semi-ambulatory telemetry systems, provided that basic channel availability and preprocessing are aligned, without the need for retraining for each small hardware variation.

#### Rodent performance and alignment with human models

On the SRD, using the same sampling frequency (256 Hz), label set (wake, NREM, REM), and TinySleepNet architecture, the model achieved a high accuracy and ROC AUC (Table 2). Although this is lower than the within-SC human performance, it shows that the same end-to-end pipeline can learn to discriminate rodent vigilance states with good accuracy.

Using an identical architecture, loss function, and training procedure for both species reduces methodological variability between human and rodent experiments. This is important for translational work because it allows differences in performance and model behaviour to be interpreted primarily in terms of species, recording conditions, and labels, rather than in terms of different model designs.

#### Cross-species transfer and translational value

The central translational result is that a model trained exclusively on SRD data retains non-trivial predictive power when applied directly to human EEG from SC via the human– rat montage. Our main methodological contribution is not a new neural network architecture per se, but the demonstration that, when constrained by an anatomically informed human– rat montage, a compact single-channel model trained solely on rat data can retain meaningful predictive value on human sleep recordings. Together with the observed cross-species metrics, which are lower than within-rat and within-SC performance but still clearly above chance (Table 2), this suggests that the cross-species mapping is capturing functionally relevant correspondences between rat cortex and human scalp, and that some of the learned signal representations are genuinely shared across species. From a translational perspective, this has several concrete implications:

- It shows that rodent datasets, which are often more controllable and can be enriched for specific genotypes or interventions, can be used not only to explore mechanisms but also to pre-train models that have measurable utility on human data.
- It indicates that some of the patterns relevant for distinguishing wake, NREM, and REM are shared between rats and humans at the level of the recorded signals, when interpreted through an appropriate montage.
- It provides a proof-of-principle that cross-species sleep staging using a common model and montage is feasible, opening a route for systematic comparison of sleep patterns across preclinical and clinical datasets.

Clinically, such a pipeline could support workflows where rodent data are used to explore candidate biomarkers or model architectures before testing on human data, thereby reducing development time and enabling more targeted evaluation in human cohorts. For example, variants of TinySleepNet or preprocessing strategies could be screened on large rodent cohorts, with the most promising configurations then evaluated, without retraining or with minimal adaptation, on human PSG. The present results show that this type of transfer yields non-trivial and practically interpretable performance. In the longer term, similar rodent-informed models could be extended to patient populations to support early detection, stratification, or treatment monitoring in sleep disorders.

### Limitations and future work

#### This work has several important limitations

First, our rodent data are limited to a SYNGAP1 haploinsufficiency model and may not capture the full range of sleep dynamics present in wild-type animals or other genetic models. On the human side, all training was performed on the SC subset of SleepEDFx, which consists of healthy adults recorded with a specific cassette-based polysomnography protocol; as a result, performance may not directly translate to other demographics, patient populations, or acquisition setups. Second, the human–rat montage is necessarily approximate and does not account for all differences in cortical geometry, folding, and volume conduction between rat cortex and human scalp. Although using a three-state wake/NREM/REM scheme reduces discrepancies between human and rodent staging, important differences in scoring conventions remain (for example, the treatment of brief arousals, transitional states, and species-specific microstructural features), which may affect cross-species comparability. Future work could refine the montage using more detailed anatomical and functional atlases and explore alternative staging schemes that better align emerging rodent substaging with human practice.

## Author contributions statement

B.C.: Conceptualisation, Methodology, Software, Validation, Formal analysis, Investigation, Writing – original draft. A.G: Data sharing. J.E. and A.G.: Supervision, Validation, Methodology, Project administration, Writing – review & editing.

All authors reviewed and approved the final manuscript.

## Acknowledgment

B.C.is funded by Epilepsy Research UK Doctoral Training Centre scholarship. This work was supported by the Engineering and Physical Sciences Research Council [grant number UKRI1659].

For the purpose of open access, the author has applied a Creative Commons Attribution (CC BY) licence to any Author Accepted Manuscript version arising from this submission.

